# BCAR: A fast and general barcode-sequence mapper for correcting sequencing errors

**DOI:** 10.64898/2026.03.27.714882

**Authors:** Bryan Andrews, Rama Ranganathan

## Abstract

**Motivation:** DNA barcodes are commonly used as a tool to distinguish genuine mutations from sequencing errors in sequencing-based assays. In the presence of indel errors, utilizing barcodes requires accurate alignment of the raw reads to distinguish genuine indels from indel errors. Existing strategies to do this generally rely on aligners built for homology comparison and do not fully utilize quality scores. We reasoned that developing an aligner purpose-built for error correction could yield higher quality barcode-sequence maps.

**Results:** Here, we present BCAR, a fast barcode-sequence mapper for correcting sequencing errors. BCAR considers all of the evidence for each base call at each position both during alignment and during final consensus generation. BCAR creates high-accuracy barcode-sequence maps from simulated reads across a broad range of error rates and read lengths, outperforming existing methods. We apply BCAR to two experimental datasets, where it generates high-quality barcode-sequence maps.

**Availability and implementation:** BCAR source code, documentation and test data are available from: https://github.com/dry-brews/BCAR

## 1 Introduction

In high-throughput sequence-function mapping, DNA barcodes are commonly appended to libraries of genetic variants. A barcode is a short non-coding sequence that occurs on the same physical piece of DNA as a variant of interest. Generally, sequencing is performed in two phases: first the entire piece of DNA is sequenced to *map* the barcodes to the putatively functional DNA variant, and second the barcode region is deeply sequenced to measure the effect of the variant, *e*.*g*., by measuring the frequency change of each barcode over the course of selection [Adamovich et al., 2022, Andrews and Fields, 2021, Jones et al., 2020, Popp et al., 2025, Starita et al., 2018]. Measuring the frequency of barcodes rather than the frequency of mutations offers a major advantage: without barcodes one cannot distinguish between a sequencing error and a bona fide mutation, but barcodes make this distinction possible. During mapping, multiple reads are collected for each barcode, with genuine mutations occurring consistently on all reads and sequencing errors occurring sporadically.

However, creating an accurate barcode-sequence map can be non-trivial when sequencing errors are common relative to bona fide mutations. *Indel* errors, where the read has a missing or extra base relative to the true sequence, are especially problematic. Once two reads fall out-of-phase, they will tend to disagree about every base call downstream of the indel (Fig. 1). Existing strategies address indels by using off-the-shelf multiple sequence alignment algorithms to align reads to a reference (e.g., alignparse [Crawford and Bloom, 2019]) or to each other (e.g., PacRAT [Yeh et al., 2022]). Then, reads with indels are filtered out (alignparse) or heuristics are used to pick which read to trust (PacRAT).

**Figure 1.**
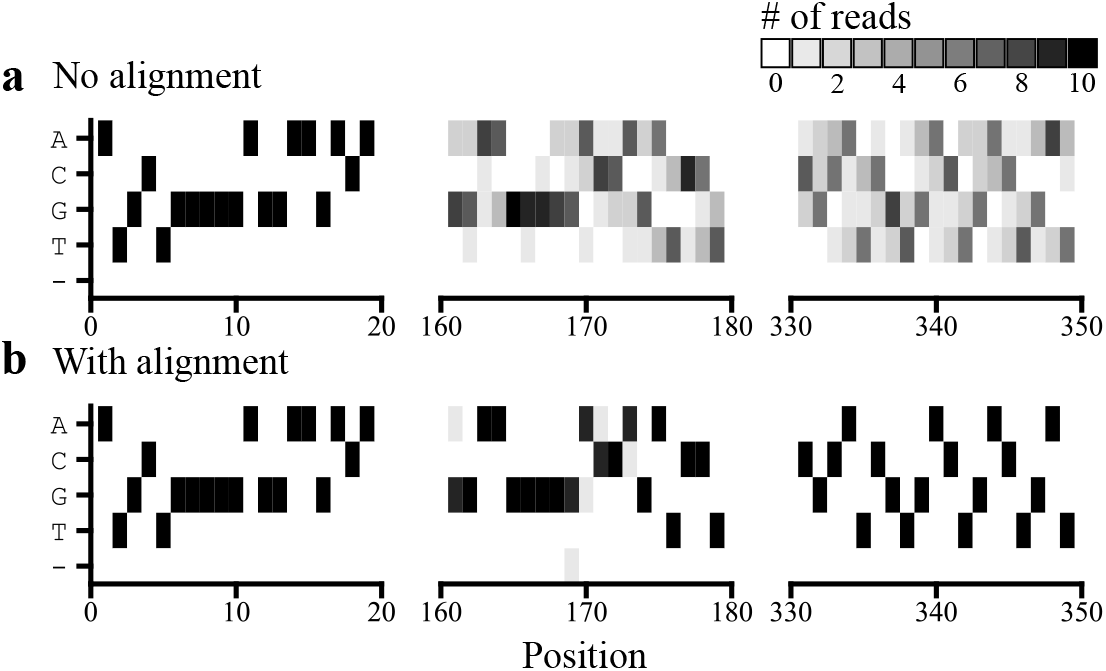
Position-weight matrices for all 10 raw reads associated with a particular barcode from an AVITI run of 16 million barcoded PDZ3 variants. **(a)** Base calls were counted at each position for all reads after adapter trimming, without alignment at the beginning (*left*), middle (*center*), and end (*right*) of the read. **(b)** As above, except the reads were aligned using Clustal Omega with default parameters [Sievers and Higgins, 2018].

Pragmatically, using filtering or heuristics to resolve indels can limit the broad applicability of an analysis tool. Fil-tering will tend to fail when the error rate is large relative to 1*/L*, where L is the read length, because most reads will have errors. Modern long-read sequencers can produce reads *>*10kb, making filtering impractical even with low error rates. Heuristics, while useful, can be highly platform-dependent, with *e*.*g*., PacRAT being explicitly optimized for Pacific Biosciences reads.

On a more principled level, commonly used multiple sequence aligners are designed for comparing homologous sequences and are not necessarily well-suited for error correction. Sequencing reads generally have associated quality scores that estimate the per-base error rate, whereas most aligners have no notion of uncertainty about a base identity. Furthermore, most aligners assume phylogeny, using a guide tree to direct alignment, but phylogeny does not apply to sequencing datasets.

Therefore, we sought to build a barcode-sequence mapper that was explicitly designed to align sequencing reads with errors. We supposed that a well-designed mapper should treat raw reads as collections of evidence about the true sequence, rather than as sequences per se, and that the mapper should handle uncertainty explicitly throughout the alignment and consensus generation process. In principle, this framework would allow us to generate accurate consensus sequences even when every read contains many errors, e.g., because the error rate is very high or because the reads are very long.

## 2 Methods

We present BCAR: Barcode Collapse by Aligning Reads. BCAR’s purpose is to quickly align the sequencing reads associated with each barcode in a dataset and generate a maximum-likelihood consensus sequence, including confidence scores for each base call. BCAR diverges from existing strategies by considering reads as matrices that contain the evidence for each base call at each position, rather than as strings of putative bases. The overall workflow consists of three parts: sorting reads by barcode, progressively aligning the reads into a single array, and converting that array into a consensus sequence with quality scores.

First, the reads are sorted on the basis of their barcode. The rationale for sorting is that the full sequencing dataset may be too large to fit in memory, and sorting the reads allows all of the reads associated with one barcode to be processed without loading the whole dataset. Therefore, the sorting algorithm itself must also avoid loading the whole dataset into memory. BCAR uses a disk-sorting algorithm that operates within fixed memory constraints by sorting small chunks of the dataset, then merge-sorting the chunks.

Second, the reads associated with each barcode are merged into a single array that contains the information from all the reads. This process is depicted in Fig. 2. Raw reads are converted to arrays where each entry represents the evidence for a base call at a given position. The arrays are globally aligned with each other, allowing for fractional match/mismatch scores when the identity of a position is ambiguous. The evidence from the aligned positions is combined, and gaps may be introduced into the combined array. This process is then repeated until all of the reads corresponding to the focal barcode are accounted for.

**Figure 2:**
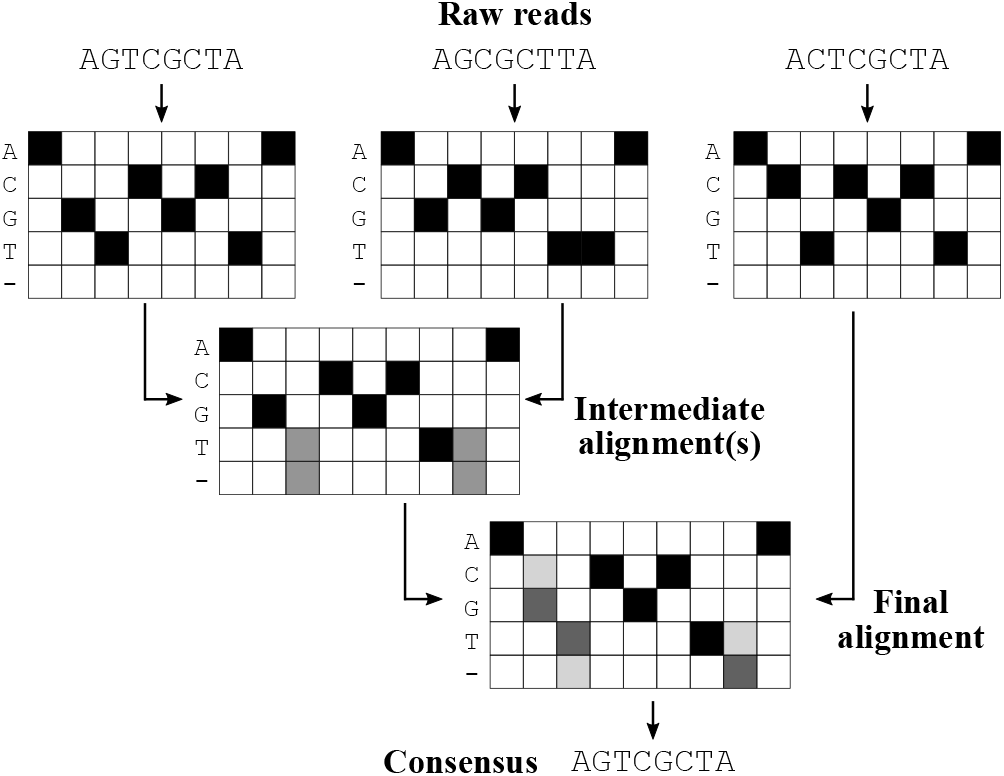
Flowchart with a toy model showing the progressive alignment strategy of BCAR. Reads are represented as arrays with entries corresponding to quality scores, then progressively aligned using a modified implementation of the Needleman-Wunsch algorithm [Needleman and Wunsch, 1970]. At each step, the new read is aligned against the current consensus, using a scaled cosine similarity to determine match scores between ambiguous positions. Gaps are removed from the final alignment during consensus generation.

Third, the final consensus array is converted back into a consensus read. At each position, the base call with the most evidence is picked, and a quality score is calculated by using Bayes’ theorem to weigh the evidence for each of the potential base calls. For more algorithmic details, see *Supplementary Material, section 2*

## 3 Results

### 3.1 Accuracy on simulated reads

To assess accuracy of reconstruction, BCAR was tested on simulated reads with defined rates of missense and indel errors. BCAR perfectly reconstructs the true sequences from simulated reads across a broad set of indel and missense rates (Fig. 3b), encompassing the measured error rates of most major sequencing platforms [Liu et al., 2024, Wenger et al., 2019, Leggett et al., 2016, Schirmer et al., 2015]. In the absence of alignment (Fig. 3a), reconstruction fails except when indels are very rare, as in Illumina reads. BCAR benefits from having more reads per barcode, but most of the benefit comes from the first ∼10 reads, and accuracy gains with more reads are modest (Fig. 3c). BCAR functions well across a broad set of realistic read lengths. It is generally easier to reconstruct shorter reads, but BCAR performance decays much more slowly than 1*/L*, as demonstrated by successful error correction on *>*100kb reads containing hundreds to thousands of errors on average (Fig. 3d).

**Figure 3:**
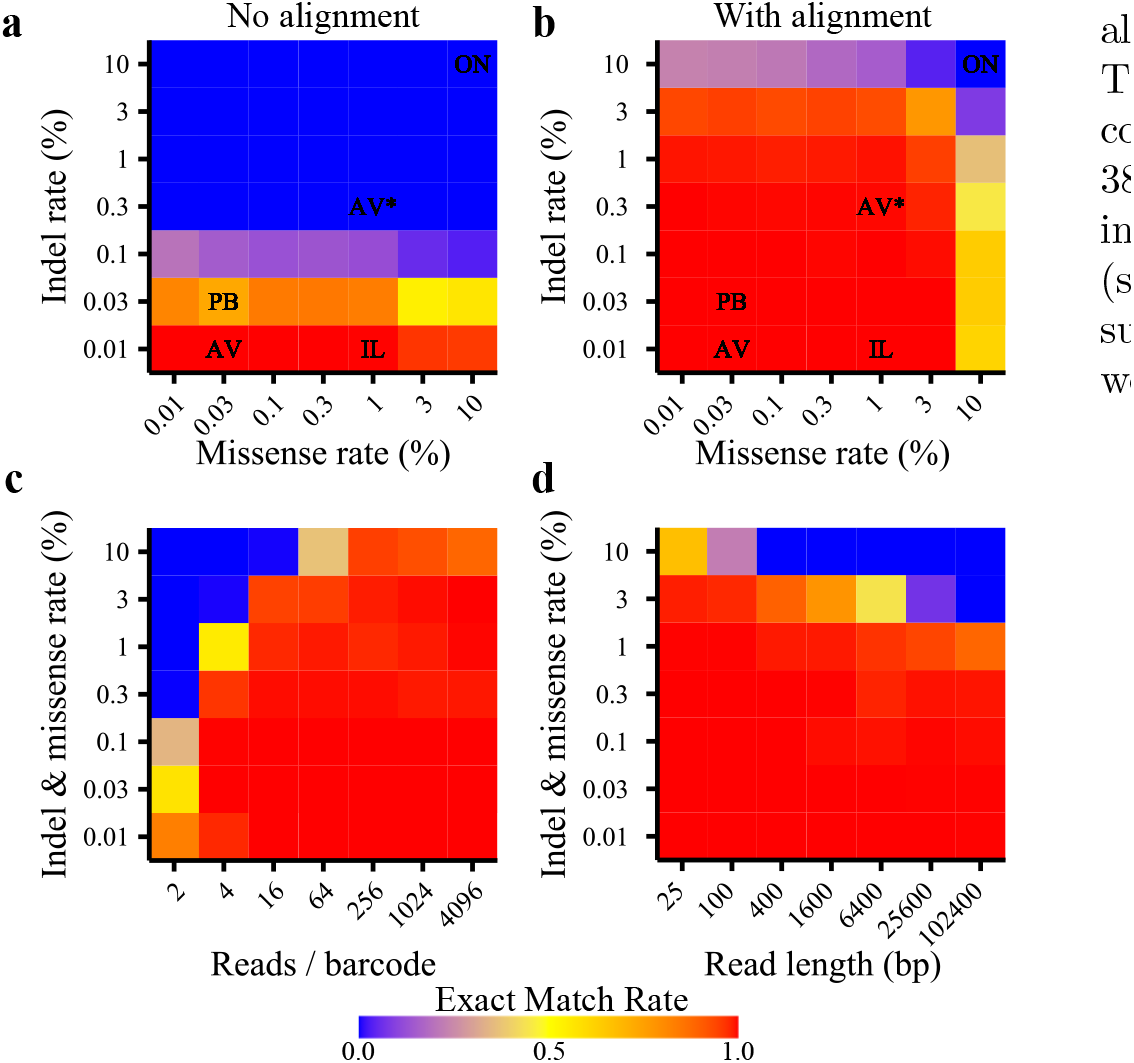
Reconstruction accuracy of BCAR on simulated reads. **(a)** Rate of reconstructing an exact match to the true sequence as a function of indel and missense error rates. Measured missense and indel rates of commercial platforms. IL: Illumina MiSeq, estimated 0.85% missense, 0.00635% indel error rate [Schirmer et al., 2015]. PB: PacBio HiFi, estimated 0.008% missense rate [Wenger et al., 2019] (0.02% this study), 0.01–0.21% indel rate [Wenger et al., 2019] (0.045% this study). ON: Oxford Nanopore, estimated 10.2% missense, 7.7% indel error rate [Leggett et al., 2016]. AV: Element AVITI, estimated 0.05% missense, 0.0075% indel error rate [Liu et al., 2024]. AV*: Element AVITI, estimated 1.6% missense, 0.3% indel error rate (this study). **(b)** The same as in (a) but without allowing BCAR to perform alignment. Most platforms have high enough indel rates to benefit from alignment. **(c)** Reconstruction accuracy as a function of error rate and reads per barcode. Having more reads is always better, but the additional benefit of more reads is mostly saturated after ∼10 reads. **(d)** Reconstruction accuracy as a function of read length. Reconstruction accuracy declines somewhat with increasing read length when error rates are very high, but with lower error rates, long reads can be reconstructed with high accuracy.

### 3.2 Application to experimental data

To assess the value of BCAR on real data, we used BCAR to construct barcode-sequence maps for two experimental datasets. The first dataset consisted of a library of 1.2 million barcoded variants of the KaiABC operon, for which we obtained 7.7 million reads of ∼3.5kb using PacBio HiFi sequencing. Using BCAR, we estimate a 0.045% indel rate per position, meaning that the average read contains *>*1.5 indels. As shown in Fig. 4, alignment is critical to obtain a high-quality consensus sequence, because when BCAR is run in no-alignment mode, the majority of consensus sequences contain low-quality base calls (Fig. 4c), which are almost entirely fixed when alignment is performed (Fig. 4d). The second dataset consisted of a library of 16 million bar-coded variants of the PDZ3 domain, for which we obtained 385 million reads of ∼350bp using Element AVITI sequencing. In this case, the benefit of alignment is less pronounced (see *Supplementary Material*), but alignment still recovers a substantial number of high-quality consensus sequences that would otherwise be low-quality.

**Figure 4:**
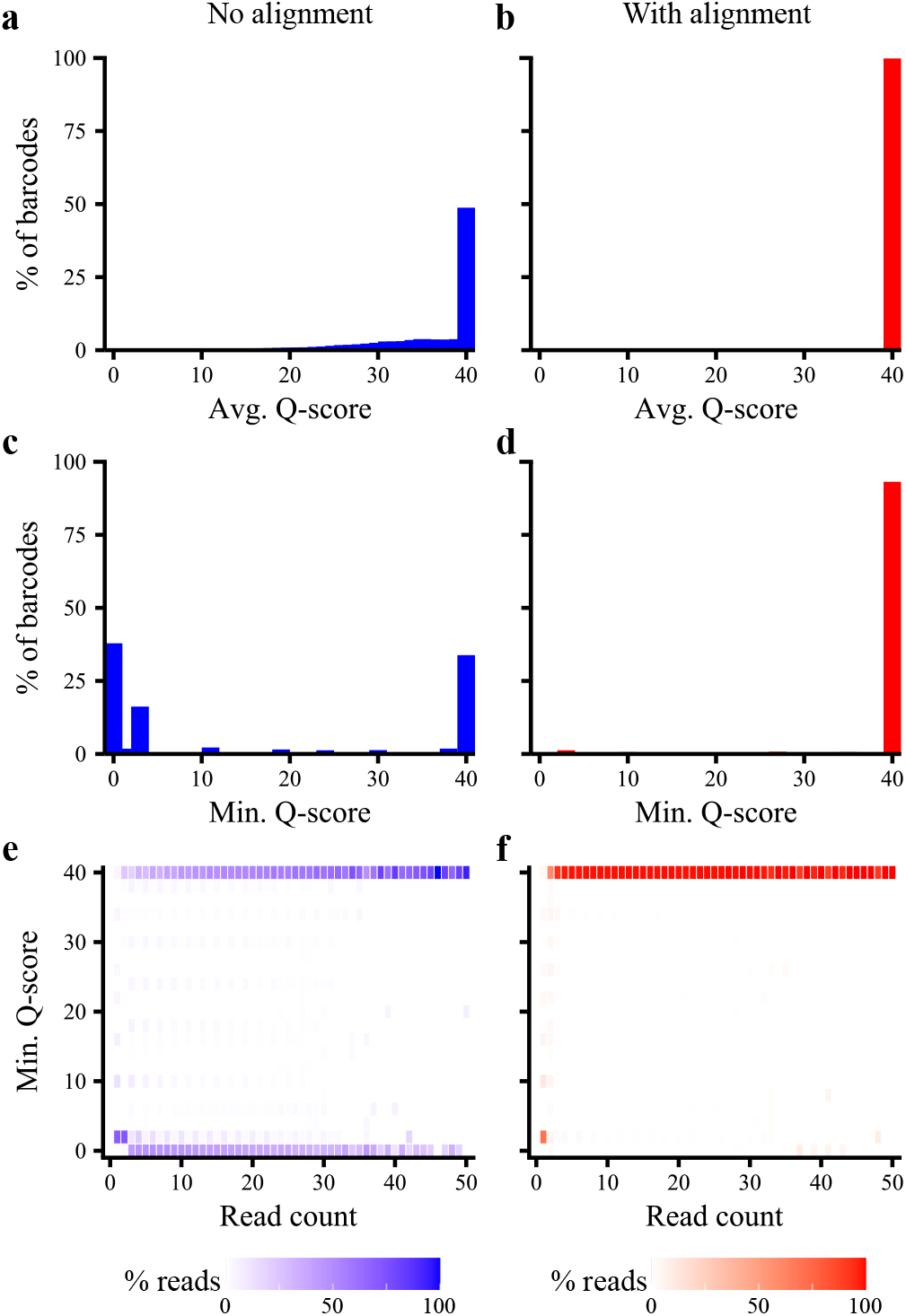
BCAR processing of PacBio HiFi reads from an experimental dataset. **(a**,**b)** Histograms of the average quality score of consensus sequences built from two or more reads, without (a) or with (b) alignment. **(c**,**d)** Histograms of the minimum quality score of consensus sequences built from two or more reads, without (c) or with (d) alignment. **(e**,**f)** Heat maps showing the minimum quality score of reads as a function of read count, without (e) or with (f) alignment. Each cell is shaded according to the percent of reads at a given read count that have the indicated minimum Q-score.

### 3.3 Comparison to existing methods

Recent progress has been made to develop tools for barcode-sequence mapping, such as PacRAT [Yeh et al., 2022] and alignparse [Crawford and Bloom, 2019], both of which focus on PacBio data and use filtering and heuristics to generate a consensus sequence for each barcode. For PacRAT, the reads associated with each barcode are aligned using MUSCLE [Edgar, 2004], which does not utilize quality scores.

Then, for positions with ambiguity, the base call from the highest-quality read is accepted as the consensus. For alignparse, the reads are pairwise aligned to a reference sequence using minimap2 [Li, 2018], and reads with indels relative to the reference are discarded.

We tested BCAR, PacRAT and alignparse for perfect reconstruction of barcoded sequences from simulated reads. Compared to existing methods, we find that BCAR is much more accurate at moderate error rates (Fig. 5a). PacRAT and alignparse both begin to fail at ∼0.1-0.3% errors, when the average read contains at least one error. BCAR, by contrast, remains high accuracy even when each read has dozens of errors. BCAR begins to fail when sequences are likely to have positions where fewer than half the reads have the correct base call (dashed line, Fig. 5a). Although BCAR can theoretically make a correct base call from a plurality of reads, in practice, positions without a majority base call are typically low-confidence and converted to “N”. Therefore, the probability of all positions having the correct base call be the majority represents a soft theoretical bound on BCAR’s accu-racy as a function of error rate. BCAR conforms very tightly to this bound when all errors are missense errors (Fig. 5a), and underperforms it slightly when errors can be missense or indels (Fig. 5b), likely due to occasional alignment errors when indels are very frequent. BCAR is also much less sensitive to read length than PacRAT or alignparse (Fig. 5c) and can achieve high accuracy with fewer reads (Fig. 5d).

**Figure 5:**
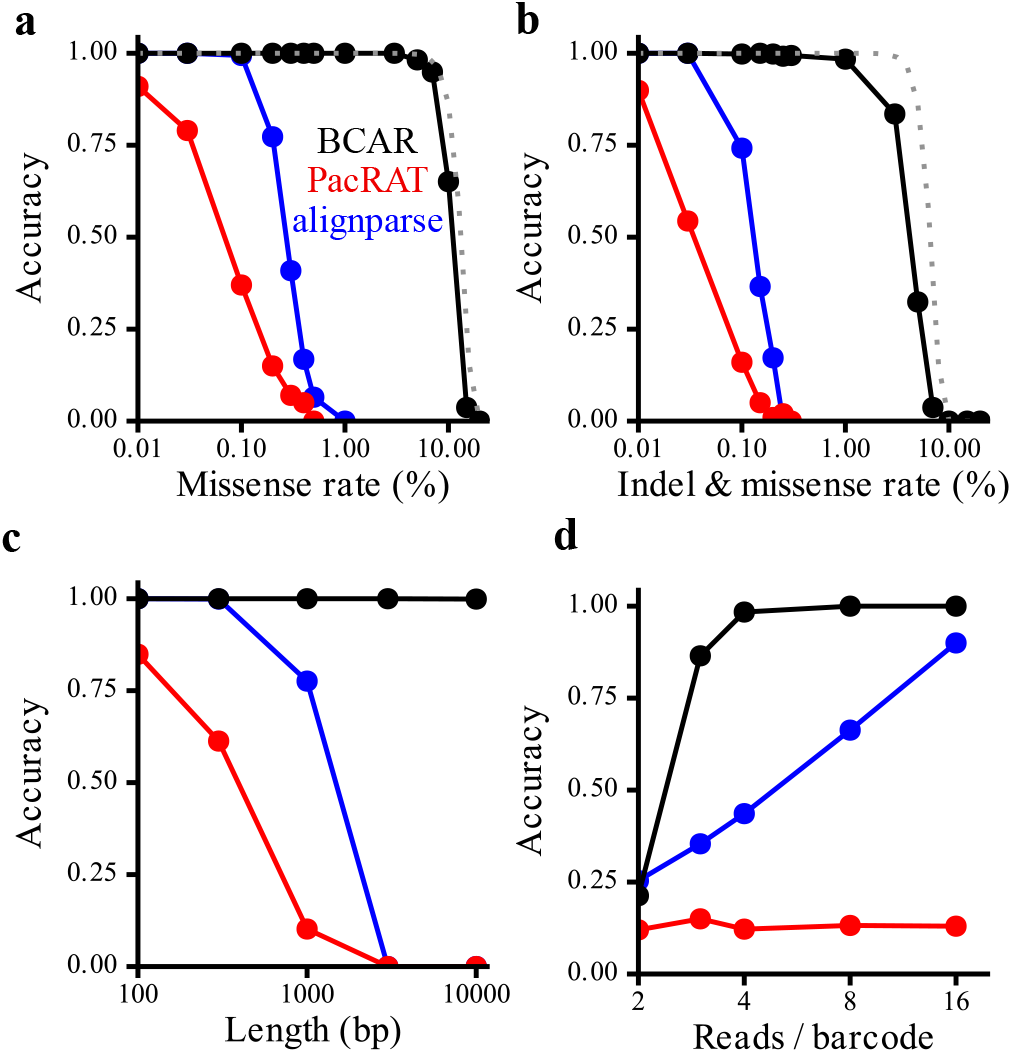
Comparison of BCAR accuracy on simulated reads to other methods. **(a)** Accuracy as a function of missense rate for BCAR, PacRAT, and alignparse. Reads were simulated at 1kb length, with 10 reads per barcode for 1000 barcodes in each condition. The barcode is not included in the sequence length. No indels were introduced. The dashed line represents the probability for a given barcode that at least half of the reads have the correct base at all positions, and serves as an approximate theoretical upper bound on the accuracy of BCAR. **(b)** As in (a), except missense and indel errors were each introduced at the indicated rates. **(c)** Accuracy as a function of the length of the target sequence. Reads were simulated with 0.1% missense and indel error rates, with 10 reads per barcode. **(d)** Accuracy as a function of the number of reads per barcode. Reads were simulated at 1kb length with 0.1% missense and indel error rates.

## 4 Discussion

Accounting for and correcting sequencing errors is a critical part of many sequencing applications. Here, we have described the development of BCAR, a fast and general tool for constructing accurate barcode-sequence maps, even in the presence of indel errors. BCAR accomplishes this by aligning the reads associated with each barcode, using a novel aligner purpose-built for handling sequencing reads. This approach mitigates some of the downsides of using existing multiple sequence aligners.

1. Multiple sequence aligners typically compare genuinely different sequences, with no notion of uncertainty about base identity. In contrast, BCAR uses quality scores to combine evidence across multiple reads, and uses a match/mismatch scoring system based on similarity to compare ambiguous positions.
2. Multiple sequence alignment is typically predicated on an assumption of phylogeny, which does not apply to sequencing reads, and utilizes a guide tree to limit propagation of alignment errors. BCAR aligns reads in order of similarity to an unaligned consensus, which serves a similar function to building a guide tree, but is more appropriate for handling sequencing errors.
3. Multiple sequence alignment is rarely applied to the scale of sequencing datasets, which can contain millions to billions of reads. BCAR is sufficiently fast and memory efficient to handle very large datasets, with submillisecond per-read processing time for typical read lengths.

Additionally, we emphasize the flexibility of BCAR. BCAR is platform agnostic rather than being designed specifically for a particular error profile or read length. Furthermore, BCAR inputs and outputs fastq files rather than relying on intermediate file formats or extensive pre-processing. BCAR does not require a reference sequence, meaning it can be applied to many experimental designs. Lastly, BCAR does not impose filtering thresholds on quality scores or read counts, but rather incorporates all available evidence in a Bayesian framework. The user can then choose how to filter their data after BCAR processing. We believe that the combination of speed and flexibility will make BCAR broadly applicable to wide variety of sequencing datasets.

## 6 Author contributions statement

B.A. and R.R. conceived the experiments, B.A. wrote the code, analysed the results, and wrote the manuscript. B.A. and R.R. reviewed the manuscript.

## 7 Acknowledgments

The authors thank Michael Rust, Diane Schnitkey and Steven Wasserman for help collecting sequencing data. The authors thank Maryn Carlson and Kabir Husain for helpful comments on the manuscript. This work is supported by funds from the National Institutes of Health grants T32AI153020 and RO1GM141697.

