## Supplementary Information for "BCAR: A fast and general barcode-sequence mapper for correcting sequencing errors"

### 1 Practical advice for running BCAR

#### 1.1 Pre-processing and post-procecessing reads

BCAR is designed to be used with other sequence processing tools. A proposed analysis pipeline is presented below. Annecdotally, we find that BCAR performs best after any pre-processing steps have already been performed (e.g., read pairing and trimming).

If indels are prevalent, it is important that the barcode occur near the start of the read. BCAR identifies the barcode purely by position, so any indels occurring prior to the barcode will cause the barcode to be read improperly. This can be easily performed using tools like CutAdapt or seqtk (https://github.com/lh3/seqtk).

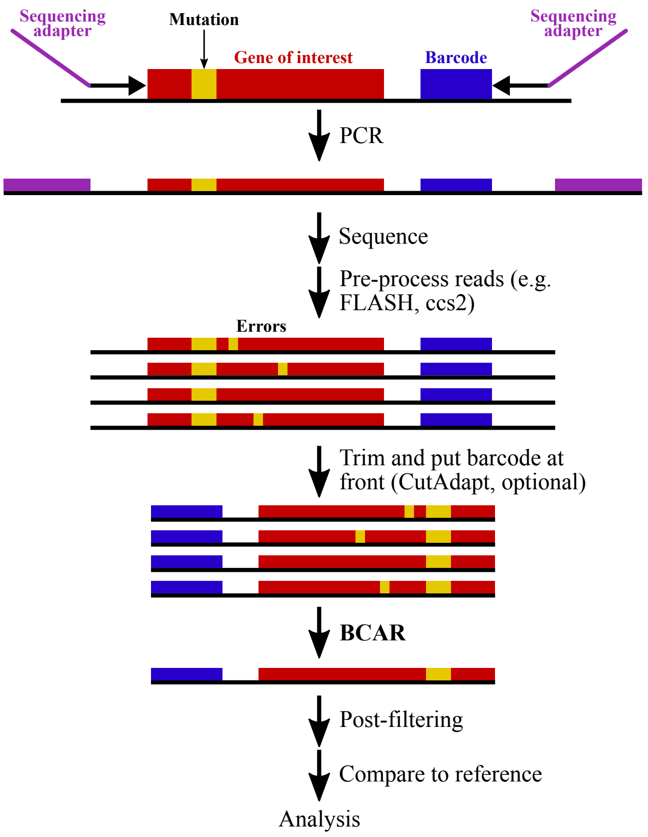


Figure S1: An example workflow that incorporates BCAR with other sequence processing tools to build a barcode-sequence map.

BCAR returns a fastq file with information about the barcode in the header, including the number of reads that contributed to it. The user may wish to post-filter their data by requiring a certain number of reads contributed to each barcode, or by thresholding the minimum or average quality score of the consensus sequence.

#### 1.2 Very long reads (>100kb)

If you have very long reads (>100kb), pay special attention to the `--max-len` flag. If you have even a single read that is longer than `--max-len`, it may cause unexpected behavior. You should either pre-trim your reads to a maximum length or determine the longest read length and set `--max-len` slightly higher than that value. For reads of this length, you are likely to run into memory constraints. Memory is shared across all active threads, so if you set fewer threads, more memory will be available to each.

##### 1.3 Paired end data

Paired-end reads are not supported by the default build, because we generally see better behavior when running all pre-processing steps (e.g., FLASH) prior to running BCAR. However, if your experimental design necessitates running BCAR on paired data, you can install a version that does handle paired-end reads by running the following commands while in the BCAR directory:

```
conda activate bcar-env
${CONDA_PREFIX}/bin/gcc -O3 -w -I${CONDA_PREFIX}/include \
-L${CONDA_PREFIX}/lib -lm -pthread -o bcar_merge_paired ./src/bc_merger_v11.c
```

Use the flags `--in-pairs` and `--out-pairs` for both the sorting and merging scripts.

##### 1.4 Very many reads (>1 billion)

In our experience, BCAR is sufficiently fast to handle hundreds of millions of reads within a few hours. If you have more reads than that and many CPUs, you can split your reads into smaller files to be processed in parallel. Suppose you sorted your dataset and determined it contains 10.5 million barcodes, and you have 10 CPUs to work with. Run the following:

```
python ./src/split_sorted_reads_into_chunks.py --in1 my_sorted_reads.fastq \
--bc-start 0 --bc-len 18 --size 1050000
```

This will split your data into 10 files with 1,050,000 barcodes that you can process separately. Afterwards, you can concatenate the BCAR output files.

##### 1.5 Many missense mutations, few indels

BCAR defaults to a gap penalty of -1.0, which is appropriate when indel errors and missense errors are approximately equally abundant. On simulated reads, BCAR is quite accurate with this gap score across a broad range of indel and missense errors. However, if you have very frequent missense errors (>1%) and quite infrequent indel errors (<0.1%), then using a more severe gap penalty may improve accuracy. A gap penalty of -3.0 is probably appropriate for many such datasets.

### 2 Detailed description of algorithm

#### 2.1 Sorting reads by barcode

Reads are sorted using a C++ implementation of a disk-sorting algorithm. Because sequencing datasets can be very large and consist of multiple files, the algorithm is designed to handle one or multiple inputs of any size in a memory-friendly way. First, the reads are loaded in 4GB chunks. For each read, the barcode is extracted based on the user-defined start position and length of the barcode. Within the chunk, reads are sorted using Quicksort [Hoare, 1962] and printed to a temporary file. When all of the input files have been processed, the temporary files are all opened and then sorted by Mergesort [Cole, 1988] into a final output file. If more than 240 temporary files are generated (∼1TB of data), then the temporary files are merged hierarchically in batches of 240.

#### 2.2 Adjusted quality scores

Raw reads consist of a string of base calls, typically A, C, G, T, or N, and a string of ASCII-encoded quality scores (also called Phred scores), where each score represents:

$$Q_{phred} = -10 * \log_{10}(p_{error}) \tag{1}$$

where $p_{error}$ is the probability that the actual base is different from the base call. Q-scores are typically scaled 1–40, though they can be higher in principle. When generating a consensus sequence, we will want to combine evidence from multiple reads. $Q_{phred}$ scores are not on an additive scale, e.g., the evidence provided by two $Q_{phred} = 20$ base calls is not equal to the evidence provided by one $Q_{phred} = 40$ base call. We define an adjusted quality score, $Q_{adj}$, which represents one-tenth the evidence conferred by a single base call with $Q_{phred} = 10$. That is:

$$p_{error} = \frac{(1/10)^{Q_{adj}/10}}{(1/10)^{Q_{adj}/10} + (9/10)^{Q_{adj}/10}} \tag{2}$$

We can convert between this score and conventional $Q_{phred}$ scores by setting the two expressions for $p_{error}$ equal to each other:

$$\frac{(1/10)^{Q_{adj}/10}}{(1/10)^{Q_{adj}/10} + (9/10)^{Q_{adj}/10}} = 10^{-Q_{phred}/10} \tag{3}$$

$$\frac{1}{1 + 9^{Q_{adj}/10}} = 10^{-Q_{phred}/10} \tag{4}$$

$$Q_{adj} = \frac{10}{\log(9)} \log\left(10^{Q_{phred}/10} - 1\right) \tag{5}$$

$Q_{adj}$ scores are on an additive scale, simplifying combining evidence from multiple reads. Furthermore, they fit naturally into a Bayesian framework for estimating base call confidence. For the 1–40 scale of $Q_{phred}$ scores, $Q_{adj}$ scores range from 0–42. BCAR considers all "N" base calls, and all base calls with $Q_{phred} \leq 3$ to contain zero evidence.

It is a quirk of Phred scores that low scores correspond to very high error probabilibities, such that a low enough Phred score actually indicate evidence *against* the putative base call. For instance, $Q_{phred} = 1$ indicates a 79% probability that the base call is incorrect. We define $Q_{adj} = 0$ for $Q_{phred} \leq 3$, rather than converting them to negative values, on the intuition that people generally do not intend to indicate negative evidence with a low score, but merely the lack of strong evidence.

In summary, we use the following conversion:

| $Q_{phred}$ | $Q_{adj}$ | $Q_{phred}$ | $Q_{adj}$ | $Q_{phred}$ | $Q_{adj}$ | $Q_{phred}$ | $Q_{adj}$ | $Q_{phred}$ | $Q_{adj}$ |
|---|---|---|---|---|---|---|---|---|---|
| 0 | 0 | 9 | 9 | 18 | 19 | 27 | 28 | 36 | 38 |
| 1 | 0 | 10 | 10 | 19 | 20 | 28 | 29 | 37 | 39 |
| 2 | 0 | 11 | 11 | 20 | 21 | 29 | 30 | 38 | 40 |
| 3 | 0 | 12 | 12 | 21 | 22 | 30 | 31 | 39 | 41 |
| 4 | 2 | 13 | 13 | 22 | 23 | 31 | 32 | 40 | 42 |
| 5 | 4 | 14 | 14 | 23 | 24 | 32 | 34 | >40 | 42 |
| 6 | 5 | 15 | 16 | 24 | 25 | 33 | 35 | | |
| 7 | 6 | 16 | 17 | 25 | 26 | 34 | 36 | | |
| 8 | 8 | 17 | 18 | 26 | 27 | 35 | 37 | | |

#### 2.3 Merging reads into an alignment

Upon observing a read, BCAR converts the read into an array, where the rows represent [A,C,G,T,-] and the columns represent positions. At each position, the cell representing the base call for the read is filled with the $Q_{adj}$ score and the other cells are filled with zero. The fifth row, "-", represents a gap character that does not appear in the raw reads but may be used during alignment.

Prior to alignment, BCAR builds an unaligned consensus of the raw reads by adding the read matrices element-wise. Then, the reads are sorted according to their similarity to the unaligned consensus, with the most similar reads being merged first to limit propagation of alignment errors. The reads are iteratively pairwise aligned using the Needleman-Wunsch algorithm [Needleman and Wunsch, 1970], adapted for ambiguous matches. While the Needleman-Wunsch algorithm typically uses fixed "match" and "mismatch" scores (commonly, 1 and -1), BCAR must compare positions where there is ambiguity about the base call. For the two vectors representing the positions being compared between the read array and the consensus array, the match score is taken to be the cosine similarity between the vectors, linearly rescaled from $[0, 1]$ to $[-1, 1]$. These values, along with a user-speicified gap penalty, are used to fill a dynamic programming array in the usual way (but see section 2.4). The traceback from this array directs which positions to merge between the two sequences. For the aligned positions, the vectors representing the positions are simply added element-wise. When a gap is proposed, the base calls from the sequence that don't support a gap are maintained, and the base calls from the sequence that does support a gap are summed and added to the gap field ("-") of the new alignment. Iterative pairwise alignment continues until all reads are accounted for.

#### 2.4 Banded alignment

To optimize speed and memory usage, BCAR uses a banded alignment strategy [Hirschberg, 1975], where the dyanamic programming array is only filled within a diagonal band of fixed width. The optimal path within the band is not guaranteed to be an optimal global alignment, but one can check whether it is optimal by comparing in to the highest possible score that would leave the band [Gibrat, 2018]. The width of the band therefore balances between speed and the likelihood of finding an optimal alignment, and the best choice of band width is a funciton of the indel rate. BCAR works by first choosing a band width as $\sqrt{L/25}$, where $L$ is the longer of the read length or the consensus length, based on guessing an indel rate of ~1% [Gibrat, 2018]. If the resulting alignment is found to be optimal, it is accepted. Otherwise, the band is progressively widened until an optimal alignemnt is found, falling back to a full global alignment if necessary.

#### 2.5 Generating a consensus

Each position in the final alignment is represented by a vector of evidence for each base call (including gaps). Positions with a plurality of evidence for a gap are skipped and do not contribute to the consensus read. For other positions, the base call with the most evidence is proposed as the true base. Evidence for all other bases (excluding gaps) is summed as evidence against the proposed base call. Bayes' theorem is applied to calculate a $Q_{phred}$ score for the proposed base call as follows. For a base call $b$ in $B \in [A, C, T, G, -]$

$$\log_{10}(p_b) = \frac{Q_b}{10} * \log_{10}(\frac{9}{10}) + \sum_{i \neq b}^{B} \left( \frac{Q_i}{10} * \log_{10}(\frac{1}{10}) \right) \quad (6)$$

So, $p_b$ represents the non-normalized probability of base $b$ being the true base at each position. The probabilities of the bases that are not proposed as the consensus base can be summed and normalized to produce a total probability of error, which can be converted to a Q-score.

$$Q_{phred} = -10 * \log_{10}\left(\frac{\sum_{i \neq max(B)}^{B} p_i}{\sum_{i}^{B} p_i}\right) \tag{7}$$

By convention, BCAR caps the quality scores $Q_{phred} \leq 40$, and for $Q_{phred}$ values of 1–3, the base call is converted to an “N”.

#### 3 Simulating reads

Barcoded reads were simulated for the purpose testing accuracy of error correction. For each barcode, a wildtype sequence was generated by randomly sampling $L$ bases with equal probability, then prepending the sequence with a 15bp barcode also randomly sampled with equal probability. Then, for each barcode, $R$ reads were generated by copying the wildtype sequence at each position while allowing for the possibility of errors. First, an indel was allowed with probability $\mu_i$, equally split between (single-base) insertions and deletions. Insertions were given a random base. Then, if no indel occurred, a miscall was allowed with probability $\mu_m$, equally split between the three mutant bases. When the true wild type base was copied, a quality score was sampled from a normal distribution, $N(\mu = 30, \sigma = 10)$, truncated to [1,40]. When a miscall occurred, a quality score was sampled from $N(\mu = 20, \sigma = 10)$, truncated to [1,40]. The barcodes were not given errors and always had a quality score of 30.

### 4 Experimental data collection.

The test data come two mutational scanning libraries. For the “AVITI” library, DNA minipreps were prepared for sequencing using a 2-round PCR strategy, which adds phased sequencing adapters on both sides, library-specific dual indices, and Truseq-style Illumina adapters. The samples were then sent to the University of Chicago genomics core for paired-end 300-cycle reads (PE300) on the Element AVITI. For the “PacBio” library, DNA minipreps were linearized by a restriction enzyme which cuts on the plasmid backbone. The linearized product was sent to SeqCenter LLC for sequencing on the PacBio Revio system.

#### 4.1 Pre-analysis read processing.

For both libraries, the raw reads were trimmed using CutAdapt to remove the adapter sequences and leave only the sequence between the start codon and the last base of the barcode [Martin, 2011]. Reads without adapters were discarded. PacBio reads can occur in either orientation, so CutAdapt was run in each orientation and the resulting trimmed sequences were put into the same orientation and combined. The reads were then all pooled and sorted on the basis of their barcodes. The AVITI dataset consists of 385 million reads covering 16 million barcodes, with $\sim$24 reads per barcode, on average. The PacBio dataset consists of 7.7 million reads covering 1.2 million barcodes, with $\sim$6 reads per barcode, on average.

#### 5 Analysis of AVITI library

The AVITI library, described in “Application to experimental data”, was analyzed by BCAR with and without alignment (Fig. SS2)

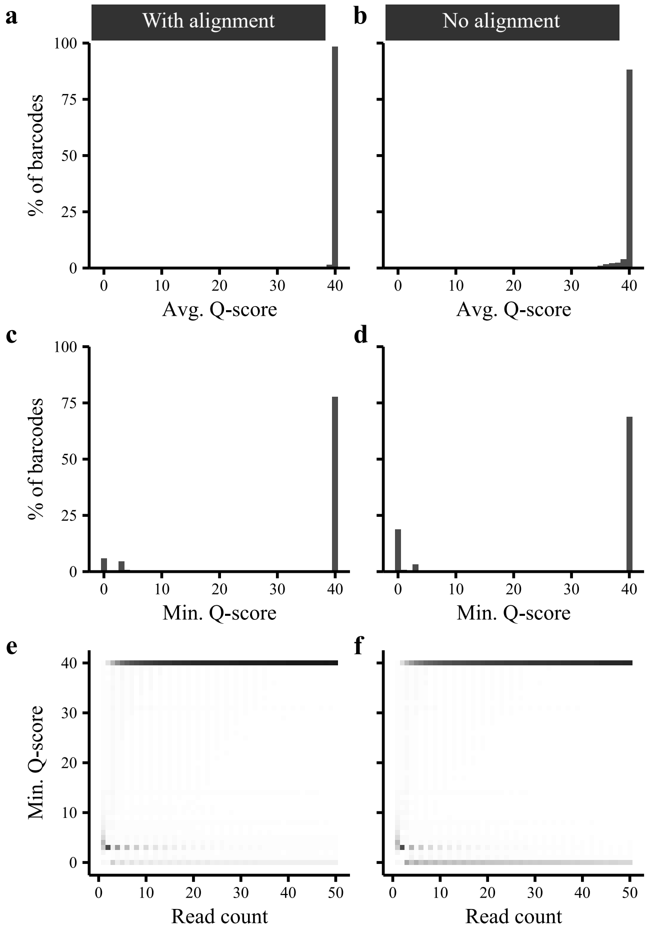


Figure S2: **(a-b)** Average Q-score distribution of consensus reads from BCAR, run with (a) and without (b) alignment. **(c-d)** Minimum Q-scores of consensus reads from BCAR, run with (c) and without (d) alignment. **(e-f)** Minimum Q-score as a function of the number of reads per barcode, for BCAR run with (e) and without (f) alignment.

Using alignment, BCAR recovers some consensus reads that would be very low quality without alignment (Fig. SS2c-d). However, the effect is less pronounced than in the PacBio dataset. This is despite the fact that, based on BCAR-estimated indel rates, the AVITI and PacBio datasets have a similar number of indels per read (1.6 indels / read for PacBio vs. 1.0 indels / read for AVITI). In addition to having slightly fewer indels per read, the AVITI dataset also have more reads per barcode than the PacBio dataset (24 vs 6). Furthermore, we observe a dramatically higher indel rate in our data compared to previously published estimates [Liu et al., 2024], and it is possible that this indicates library preparation artifacts such as PCR strand-exchange, rather than true sequencing errors. BCAR may be less able to correct these artifacts, as it designed to correct random, rather than systematic, errors.

### 6 Speed

Sorting a fastq file is $O(N * \log(N))$, and is generally IO-limited. The speed of merging reads is more complicated, depending on the length of reads and the frequency of indels. Because of the banded alignment strategy, in the best case, merging reads is $O(N * L)$, wherease in the worst case, it is $O(N * L^2)$. In simulations, we see $O(N * L^k)$, where $k \approx 1.25$ for indel rates of $10^{-4}$ and $k \approx 1.85$ for indel rates of $10^{-1}$ (Fig. SS3). Average memory usage mirrors speed, but peak memory usage should be assumed to be $O(N * L^2)$.

Datasets with many short reads will require more time to sort, whereas datasets with fewer long reads will require more time to merge. For example, the PacBio dataset with 7.7 million ~3500bp reads took only ~4 CPU-minutes to sort, but ~43 CPU-minutes to merge. In contrast, the AVITI dataset with 385 million ~350bp reads took ~8 CPU-hours to sort, but only ~3 CPU-hours to merge. Both tests were performed on 24–thread compute nodes (Intel E5-2680 v4 2.40 GHz).

Because merging operates on a sorted dataset, it can operate faster by reading over different parts of the dataset at the same time (*e.g.,* using different CPUs). We do not implement this strategy natively in case it could cause disk thrashing or other undesired behavior on different systems, but see section 1.4 for breaking sorted reads into separate files.

#### 6.1 Speed benchmarks

We tested the speed to bcar_merge on simulated reads with varied length and reads per barcode. Testing was performed on a 24–thread compute node (Intel E5-2680 v4 2.40 GHz). Except where otherwise noted, we used 10 reads per barcode, a 1kb read length prepended by a 15bp barcode, a 1% missense error rate, and a 1% indel error rate.

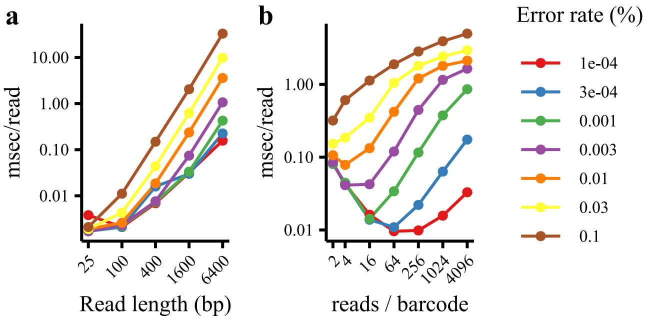


Figure S3: **(a)** Time taken per read as a function of read length. Note the log scale on both axes. At low error rates, time scales approximately linearly with read length (slope=1 in log space), whereas at high error rates, time scales closer to quadratically (slope = 2). **(b)** Time taken per read as a function of the number of reads per barcode. Increasing error rate increases the time takes per read, likely due to the increased probability of needing to widen the banded alignment.
